# What is pathogenicity? Structural vulnerabilities lead to distinct phenotypes in a membrane protein, ATP1A3

**DOI:** 10.64898/2026.10.13.758912

**Authors:** Kathleen J. Sweadner, Laurie J. Ozelius, Aloysius Domingo, Kotchaphorn Mangkalaphiban, Ihtsham Haq, Allison Brashear, Elena Arystarkhova

**Affiliations:** Department of Neurosurgery, Massachusetts General Hospital, Boston, MA, USA; Harvard Medical School, Boston, MA, USA; Department of Neurology, Massachusetts General Hospital, Boston, MA, USA; Department of Neurology, University of Miami Miller School of Medicine, Miami, FL, USA; Jacobs School of Medicine and Biomedical Sciences University at Buffalo, Buffalo, NY, USA

## Abstract

Distinct protein features of pathogenic variants are illustrated here for a gene with an extraordinarily wide range of neurological syndromes: ATP1A3, the neuronal Na,K-ATPase. Vulnerable structure features produce mechanistically different consequences, from loss or gain of function to maladaptive cell biological responses. Variants in specific structural elements (glycines, prolines, and β-sheets) link severe phenotype to impaired folding during biosynthesis. In contrast, impaired protein-protein interactions may result in brain malformation. Defective RNA transcripts, on the other hand, exhibit a range of benign to severe phenotypes, likely due to efficacy variability in the defensive degradation of aberrant mRNA and protein. Variant interpretation will benefit from understanding structure-specific vulnerabilities.

## Introduction

Na,K-ATPase (the sodium pump) is a critical membrane protein^1^. The catalytic (α) subunit has four separate domains (substructures) that coordinate movements to transport ions uphill. The protein folds in the endoplasmic reticulum membrane^2^. ATP1A3 has a Z score for missense variants of 10 (gnomAD 4.1), indicating strong selective pressure^3^, yet we currently know of high confidence pathogenic variants in only 88 amino acids out of 1013.

Na,K-ATPase catalyzes the energy-dependent exchange of three intracellular sodium ions for two extracellular potassium ions, and it is critically important in the brain to support and regulate electrophysiological activity. Three α subunit paralogs are associated with genetic disorders. Each paralog causes both milder and more severe phenotypes, but none is more diverse than ATP1A3, which is expressed in neurons. Eight syndromes associated with ATP1A3 have well-characterized phenotypes. In order of discovery, there is rapid-onset dystonia-parkinsonism (RDP)^4^; alternating hemiplegia of childhood (AHC)^5,6^; cerebellar ataxia, pes cavus, optic nerve atrophy, and sensorineural deafness (CAPOS)^7^; developmental and epileptic encephalopathy (DEE; DEE99 for ATP1A3)^8^; relapsing encephalopathy with cerebellar ataxia/ fever-induced weakness and encephalopathy (RECA/FIPWE)^9,10^; neonatal onset of dystonia with dysmorphism of the face, encephalopathy with developmental delay and cerebellar hypoplasia without hemiplegia (D-DEMØ)^11^; spasticity presenting like cerebral palsy^12^; and polymicrogyria (PMG)^13-18^, a malformation of cortex layer organization and gyrification. Remarkably, three of these disorders change single amino acids: CAPOS (p.Glu818Lys), RECA/FIPWE (p.Arg756His, Cys, or Leu), and cerebral palsy-like spasticity (p.Pro775Leu).

Concrete evidence of different mechanisms of pathogenicity in ATP1A3 has been established by laboratory experiments on selected variants. Some RDP variants are inactive, and others have altered affinities for Na^+^ and K^+^, resulting in reduced activity at physiological ion concentrations^19^. This produces altered membrane potentials^20^ and increased cytoplasmic Na^+ 21,22^. In oocytes certain AHC variants had a dominant negative effect on pump current, indicating a gain of function^23^ as well as loss of function^24^. Two DEE variants exhibited protein misfolding and activation of the Unfolded Protein Response (UPR), including endoplasmic reticulum (ER) retention and pro-apoptotic signaling^25,26^. There are PMG variants that may alter the interaction between α and its β subunit^13^. For the three single-residue pathogenic variants, each had different properties: the CAPOS variant showed altered ion affinities and a weaker voltage dependence^27,28^. The RECA/FIPWE variant, always triggered by fever, showed normal expression but inactivation and temperature-sensitive degradation in the febrile range^29^. The spasticity variant showed both inactivation and an inward leak current of Na^+12^. Each single-residue phenotype has been reported in between eight to eighty patients.

Here the investigation of variants in protein structures uncovered principles that can be applied broadly to genetic variants. There are distinct pathogenicity mechanisms, and multiple structural points of vulnerability. **Box 1** illustrates some important concepts. The structure building blocks, α-helixes and β-sheets, are stabilized by regular internal hydrogen bonding. Their side chains face outward, so most substitutions act more on the packing of α-helices or β-sheets against neighboring structures than on the structure itself, facilitating long-range consequences (**Box 1a**,**b**). β-sheets assemble from separated strands that must first align to nucleate, and a bulkier variant in a strand can prevent the sheet from forming at all (**Box 1b**). Glycine has no side chain, which allows the backbone to make tight turns and lets helices pack closely; any substitution that introduces a side chain can interfere (**Box 1c**). Proline has restricted rotation and so it breaks helices; either the loss or gain of a proline can be damaging (**Box 1d**). In the final structure, a single residue may hold several distant segments together, as an arginine does through hydrogen bonds to four other segments; its substitution causes temperature-sensitivity^29^.

### BOX 1

**Vulnerable features of protein structure**

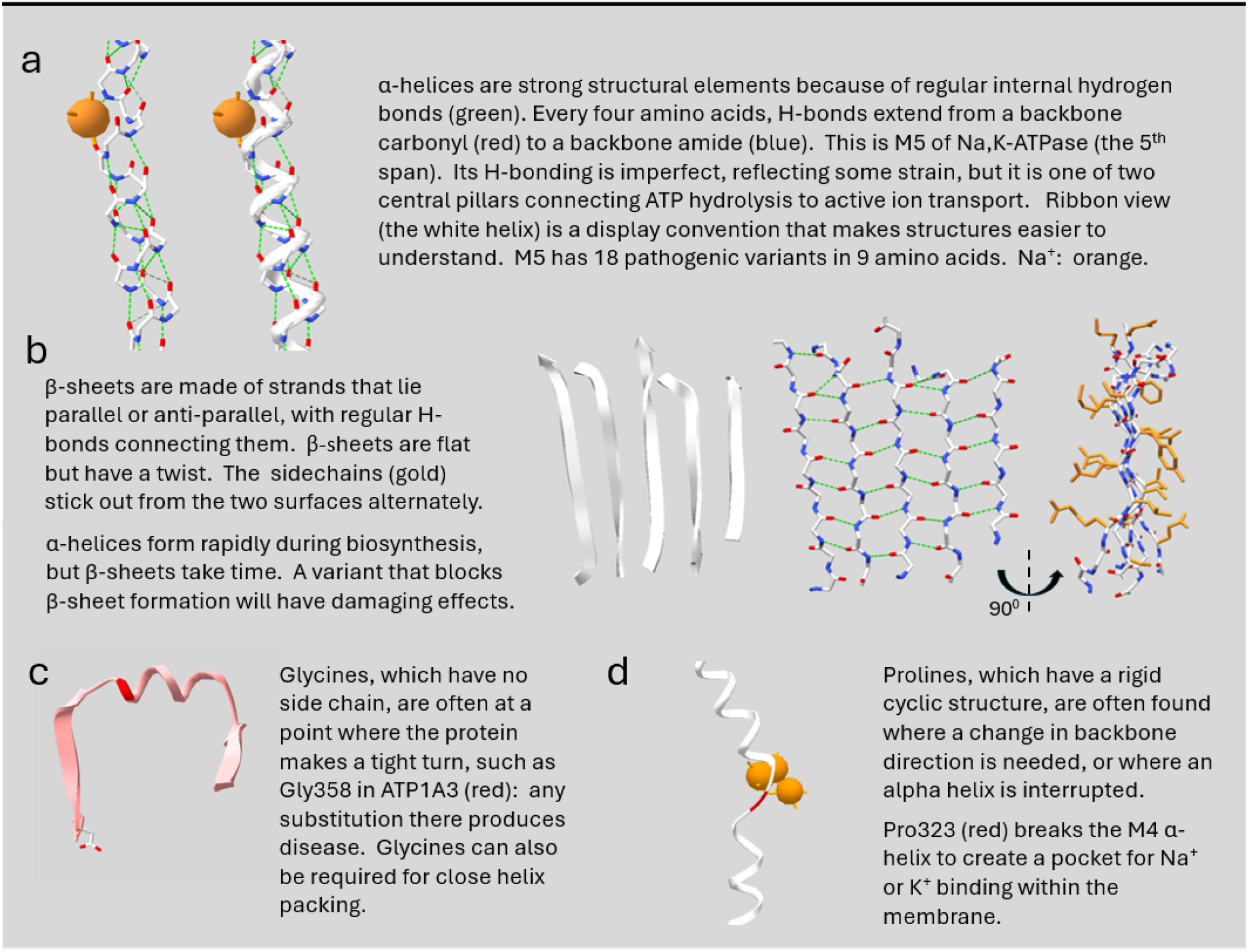

### Strategy

Many clinicians consider ATP1A3 symptoms to be on a continuum because of patients with intermediate symptoms and uncommon manifestations^30^. Our question was whether useful general principles can nonetheless be derived from structure-mechanism-phenotype relationships. We currently know of 274 different variants in ATP1A3, including alternate amino acid substitutions in single codons, from our ongoing study, publications, meeting abstracts, public databases, patient registries, social media, and discussions with physicians. All had symptoms on the spectrum for ATP1A3. We investigated variants from four major phenotypes of ATP1A3-associated disease: RDP, AHC, DEE, and PMG.

We focused on variants in the 88 amino acids that have laboratory validation or that meet these criteria: RDP variants were included if they were familial, recurred independently, or if there were two or more substitutions of the same amino acid. RDP singletons were included if they were in our consented RDP cohort, published in a case report, or if the corresponding variant in ATP1A2 produced familial hemiplegic migraine^31^. We required variants for AHC without experimental validation to be reported independently at least twice because single reports of novel variants could be false positives. We also excluded variants with intermediate phenotypes between RDP and AHC or that manifested as RDP or AHC in different individuals. This is because mixed phenotype results could be due to structural alterations that have two deleterious outcomes (for example changes in ion affinity as well as in ion conductance), and these might obscure phenotype-structure relationships. There are a limited number of severe cases with or without PMG, and all were included. They overlap in neurological function and developmental delay, but examining PMG variants separately proved to be crucially informative.

## Results

### Loss of pump activity is not a predictor of phenotype

The differences in ATP1A3 patient syndromes are not due to how much ion pump activity is lost. Because Na,K-ATPase activity is required for cell survival, a screen for pathogenicity is to test cell viability by exogenously expressing inhibitor-resistant ATP1A3 variants and inhibiting endogenous ATP1A1 with ouabain. If the exogenous ATP1A3 has enough residual activity, the cells will live. Representative variants for RDP, AHC, RECA/FIPWE, DEE, and PMG have all produced either survival or cell death (**Table 1**).

**Table 1.** Phenotype is not a function of loss of Na, K-ATPase transport activity. Ouabain-resistant ATP1A3 cDNAs were mutated and expressed in cells. The endogenous ouabain-sensitive ATP1A1 was inhibited with a low concentration of ouabain. In the absence of any exogenous ATP1A3, cells die in 24-36 hours, and in the presence of wild type ATP1A3 they survive. The table shows whether or not each mutation had enough residual activity to keep the cells alive. There was no correlation with patient phenotype. † These data were published in Vetro et al. 2021. * PMG variants.

| child or adult |  |  | child (fevers) |  |  | infant or toddler |  |  | in utero |  |  |
| --- | --- | --- | --- | --- | --- | --- | --- | --- | --- | --- | --- |
| RDP |  | cells survive | RECA/FIPWE CAPOS |  | cells survive | AHC |  | cells survive | DEE, PMG |  | cells survive |
|  | no | yes |  | no | yes |  | no | yes |  | no | yes |
| E277K |  | ✓ | R756H |  | ✓ | G755S | ✓ |  | L292R<sup>†*</sup> | ✓ |  |
| F297del |  | ✓ | R756C | ± | ± | D801N | ✓ |  | G316V<sup>†</sup> | ✓ |  |
| D366H | ✓ |  | R756L | ✓ |  | S811P | ✓ |  | D742Y | ✓ |  |
| T613M |  | ✓ | E818K |  | ✓ | E815K | ✓ |  | D887Y<sup>†*</sup> |  | ✓ |
| D743H | ✓ |  |  |  |  | L839P |  | ✓ | L888P<sup>†*</sup> | ✓ |  |
| I758S | ✓ |  |  |  |  | C927W |  | ✓ | P972del<sup>†</sup> |  | ✓ |
| D923N |  | ✓ |  |  |  | G947R | ✓ |  | L924P<sup>*</sup> | ✓ |  |

### Structure distributions of clinical variants in RDP, AHC, and DEE

All clinical variants are in the α (catalytic) subunit of Na,K-ATPase. It is a 10-span, multi-domain protein that achieves active transport by the motion of three cytoplasmic domains (A, N, and P) driven by the hydrolysis of ATP (**Fig. 1a**)^32,33^. The “stalk” has important connections between the membrane domain and the mobile domains. The motions of the A and N domains bend and tilt the P domain, which moves transmembrane spans relative to each other analogous to pistons, exposing the central ion binding site alternately to the extracellular and intracellular surfaces. The openings alternate without making a continuous pathway^34^. The β subunit (and small FXYD subunit, not shown) each contribute one membrane span, but they are not mechanistically involved in ion transport and have no relevant variants.

**Fig. 1.**
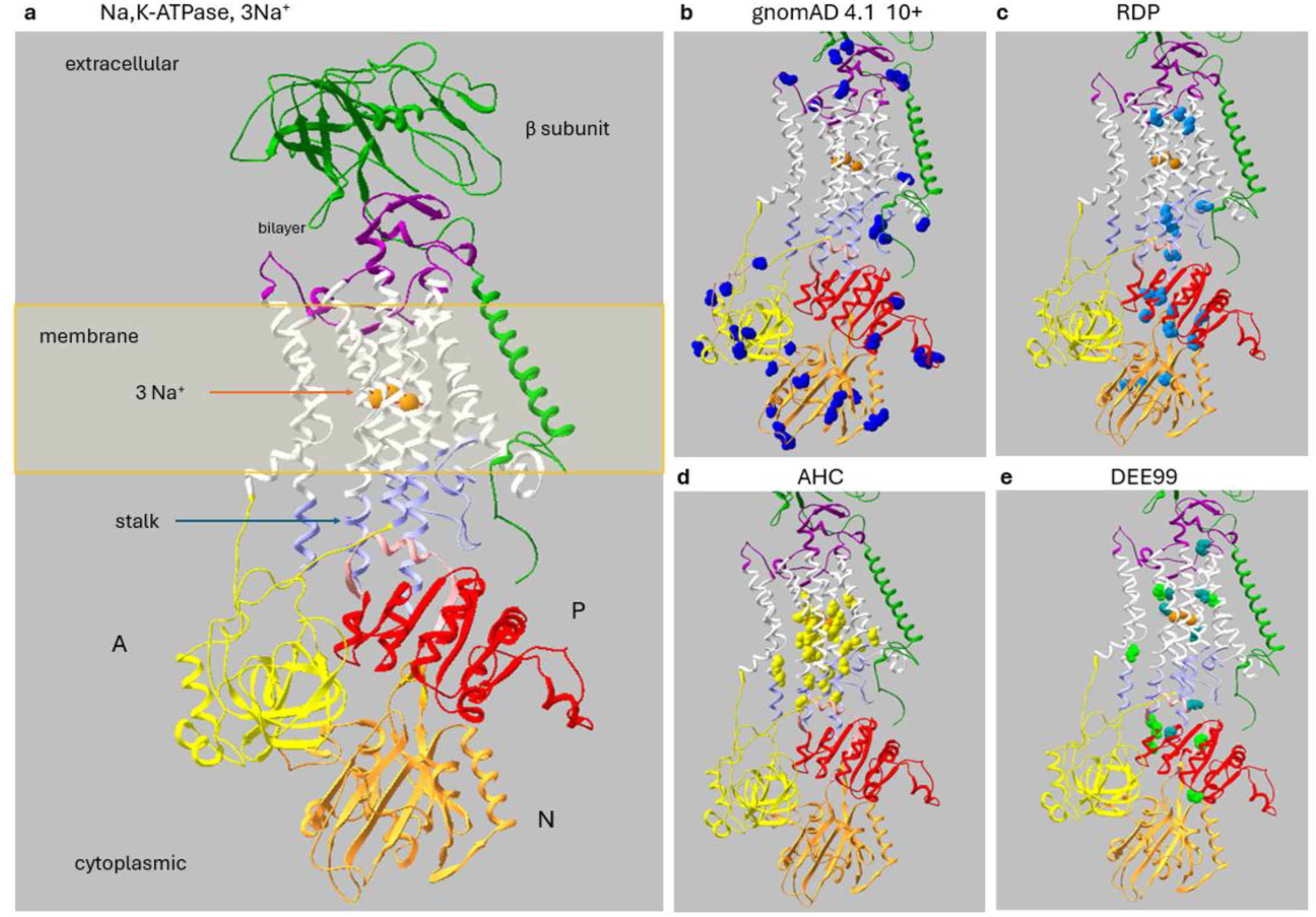
Structure distribution differences between benign and pathogenic variants. **a**, Na,K-ATPase α and β subunit structure. The cytoplasmic domains are named N for nucleotide binding, P for phosphorylation, and A for actuator. The small FXYD modulatory subunit is hidden for clarity. **b**, The most abundant gnomAD 4.1 variants that should be benign (ten or more cases in each of 38 amino acids) (blue) are represented by their three backbone atoms in spacefill. Variants in two amino acids in the N-terminus are not visible because the first 22 amino acids are unstructured and not resolved; they would be adjacent to the yellow A domain. The four benign residues in the P domain are in an extension (the hook) that is not part of the central β-sheet. **c**, Variants in cases with the mildest syndrome, RDP (teal blue). They are found far from the ion binding site, and three are in the N domain. **d**, Variants in cases with AHC, alternating hemiplegia of childhood, and one with spasticity, are all in the cytoplasmic half of the ion pathway and the stalk (yellow). **e**, Variants in cases with developmental epileptic encephalopathy, DEE (green and teal). The PDB file used is 3WGU.

As proxies for benign mutations, we used variants found in multiple individuals in the database gnomAD 4.1^35^. Altogether there were 138 gnomAD missense variants (including alternative substitutions) in 114 amino acids. Of those, variants in 38 amino acids occurred in 10 or more individuals (**Fig. 1b**). These common variants were absent from the transmembrane spans and all but a distal part of the P domain.

The distributions of disease-causing variants are not random at all. RDP variants are found in the P domain, the N domain, and peripherally in the transmembrane domain in **Fig. 1c**. Note that they are distant from the three sodium ions in the central ion-binding pocket. Nonetheless it has been demonstrated that many reduce ion affinity. Importantly, it has been shown that this is caused by long range effects: an alteration in one place can propagate to affect ion affinity by its effect on the mutual interactions of the central α-helices or through H-bond networks.^19,36^ There is no evidence that rules out haploinsufficiency as the fundamental mechanism for dominance in RDP.

AHC variants that occurred at least twice are in the transmembrane spans and the stalk (**Fig. 1d**), a very different distribution^37,38^. All amino acids that directly surround the ions have variants that cause AHC. Notice, however, that AHC variants are absent from the extracellular half of the membrane domain. This suggests that they have a gain of function property impacting only the central and cytoplasmic part of the ion pathway, where Na^+^ enters and K^+^ exits, and where protons participate in the neutralization of charged amino acids^39^.

Intractable infantile epilepsy with severe developmental delay (DEE, DEE99 = ATP1A3) is the most-reported severe manifestation (**Fig. 1e**). Many severe cases have grossly normal MRIs, although microcephaly and atrophy can occur. These variants cluster in the membrane domain or the P domain. Variants are distributed evenly across the membrane domain. What is notable is that all seven of the variants colored teal blue either change a glycine to something else or lose or gain a proline. Glycine has no side chain, which lets the backbone make tight turns and lets helices pack closely against each other; any replacement adds a side chain that gets in the way. Proline is bulky and stiff, so removing one can prevent a turn or a helix break that the structure depends on, while adding one elsewhere can break a helix that was meant to be whole. These changes can have major impacts on the protein’s ability to fold into the correct structure during biosynthesis (**Box 1**). We previously documented protein misfolding in cells expressing p.Leu924Pro, a variant found in multiple individuals with DEE symptoms^25,26^. In **Fig. 1e** the transmembrane variants are in crucial bends or in α helices. In contrast, the DEE variants in the P domain have distinctive features that will be discussed below.

It is notable that the A and N domains have many benign variants, but few qualifying pathogenic variants. The paucity may reflect reduced detection because of milder or delayed phenotypes. These two domains move dramatically during ion transport. The N domain binds ATP and brings it to the active site to transfer phosphate to an aspartate. Then the A domain rotates in to hydrolyze the phospho-aspartate. One individual with a variant in the aspartate has RDP^26^, consistent with loss of activity alone.

### Vulnerability of the ATPase P domain β-sheet

The P domain consists of a central β-sheet surrounded by α-helices. **Fig. 2a and b** illustrate a striking structural difference for mild and severe pathogenicity: all of its RDP variants (delayed onset) are in the surrounding α-helices, and its DEE variants (manifesting at birth) are in the central β-sheet.

**Fig. 2.**
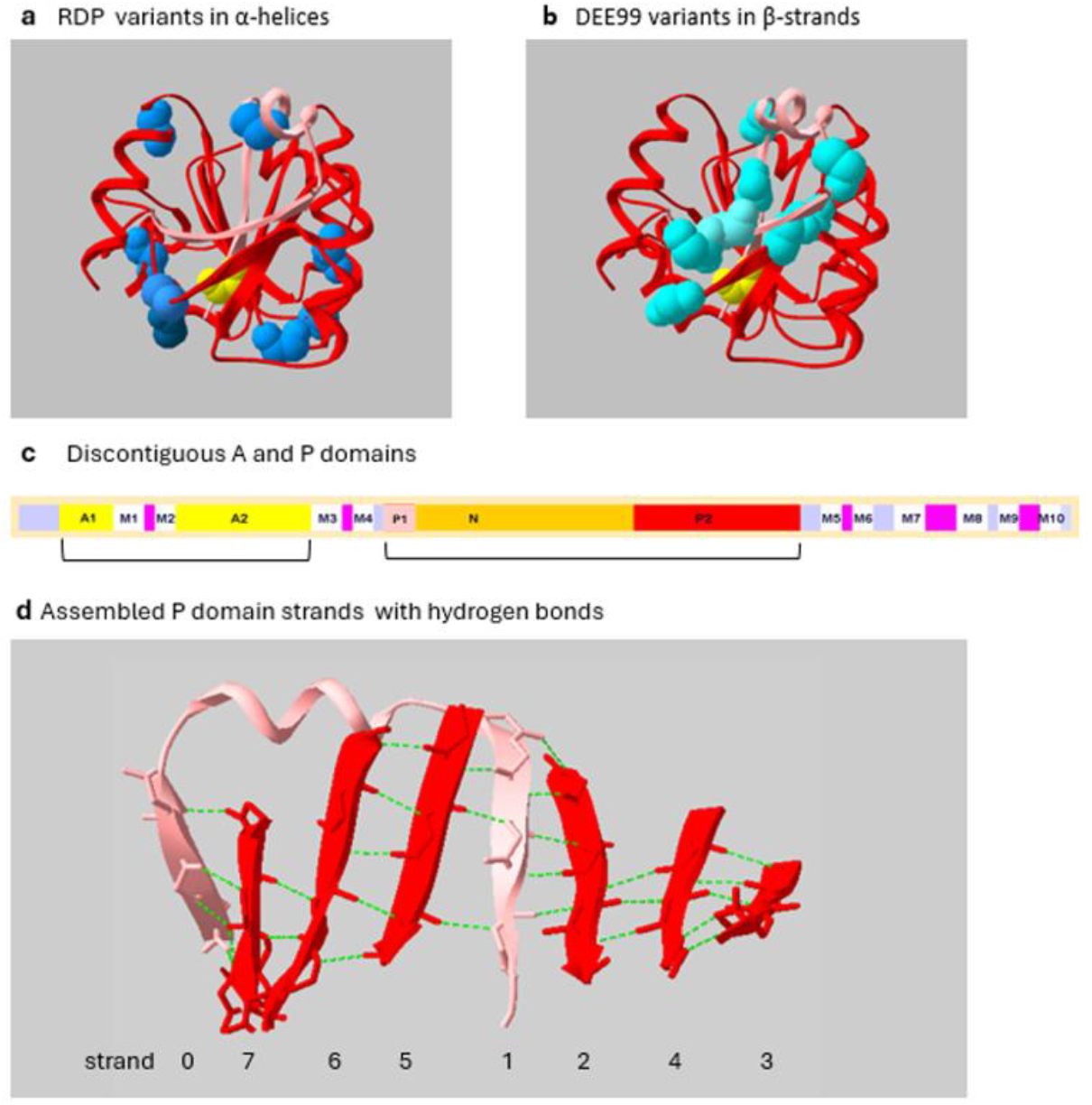
Mild vs. severe variants: α-helices vs. β sheet. **a**, RDP variants in the P domain are in the α-helices surrounding its central β sheet. The yellow atoms mark the aspartate residue that is covalently phosphorylated during the transport cycle. **b**, DEE variants in the P domain are in the central β sheet. **c**, Diagram of the ATP1A3 transcript mRNA. The A domain (yellow) and the P domain (pink and red) are both discontiguous and must assemble after translation. **d**, The P domain’s β sheet has two halves, P1 and P2, that interdigitate after the N domain is translated. The first strand is numbered zero because it is the only one that is anti-parallel in its orientation. Strand 1 from P1 has to insert between strands 2 and 5 of P2. All of the variants in **b** are bulkier and likely to interfere with β-sheet assembly.

The P domain, which is highly conserved in all P-type ATPases, has a structure that is challenging for biosynthesis. The linear arrangement of Na,K-ATPase domains on the mRNA transcript is shown in **Fig. 2c**. There are five transmembrane hairpins and their connecting loops. The A domain is interrupted by a transmembrane hairpin, and its two parts, A1 (helices and coil) and A2 (β-sheet), bind to form a stable complex. The P domain is interrupted by the entire N domain. During biosynthesis P-domain β-strands emerge from the ribosome in the numerical order shown (**Fig. 2d**) but strands 0 and 1 (pink) are in P1, and they have no strands to associate with until after the N domain is translated and translation of P2 begins. If a bulkier DEE side chain substitution in a β strand prevented the interdigitation of P1 and P2 β strands, it should suspend folding (**Box 1b**) and elicit the integrated stress response (ISR); here the substitutions are all bulkier. The ISR entails transiently inhibiting normal protein synthesis and activating a complex program of defensive responses. This disrupts normal cell function, and if it is not successful leads to apoptosis^40^. In contrast, none of the RDP variants are likely to impair folding much since the α-helices are at the P domain’s surface, which is largely aqueous except for transient domain-domain contacts. Like the effects of glycine and proline variants, the logical conclusion is that seriously impaired folding produces severe neurodevelopmental symptoms associated with DEE while the local-only effects of α-helix side chain alterations fit the milder phenotype seen in RDP.

### Variants that cause brain malformation

The most damaging ATP1A3 variants cause polymicrogyria, a brain malformation that develops in utero. Infants with PMG exhibit symptoms that overlap extensively with DEE, but the consequences are more profound. In **Fig. 3a**, 18 cases have occurred as 13 missense PMG variants in 12 amino acids shown on the structure. One loses a glycine and six gain a proline. Four (red) are in the membrane domain and three (aqua) are in the P domain, like DEE variants. In the P domain, the three PMG variants could all interfere with interdigitation of the P1 and P2 β-sheet (**Fig. 3a**,**b**), as could K717del (**Fig. 3b**)^15,41^. The overlap with DEE phenotype is logical: DEE cases in other genes are caused by aberrant neuronal migration causing positional mismatch of excitatory and inhibitory neurons and intractable epilepsy^42,43^. Such cortical dysplasia is seldom obvious in MRIs but can be detected in specimens, currently scarce for ATP1A3 cases^17^.

**Fig. 3.**
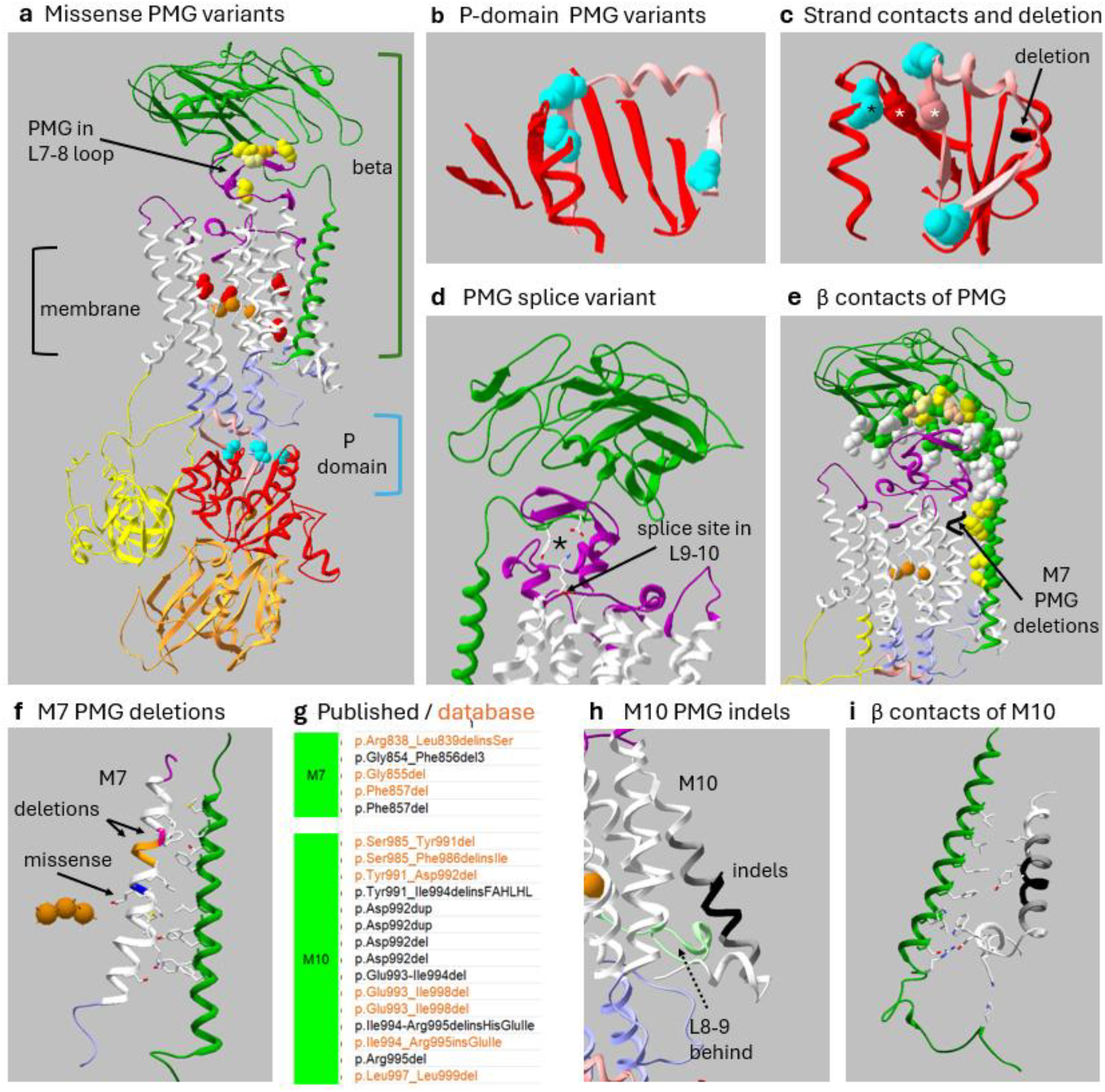
Classes of PMG variants. **a**, There are three distinct groups of missense PMG variants: those interacting with β subunit (shades of yellow), those buried in the membrane (red), and those in the P domain (aqua). **b**, The three PMG variants in the P domain should interfere with the interdigitation of the P1 (pink) and P2 (red) β strands. **c**, Rotated, showing the variant in an α-helix (black asterisk) could impair sheet assembly by displacing amino acids in P1 and P2 (white asterisks). Another α-helix has a single amino acid PMG deletion (black). All are likely to disrupt P domain folding. **d**, Splice site variant at K764, normally close to two residues in β (asterisk). Any cryptic splice that alters the L9-10 loop would disrupt the interaction. **e**, β subunit amino acids that contact the α subunit. Yellow backbone atoms mark the amino acids contacted by PMG variants, while white marks the rest. **f**, All interacting residues in M7 and the β span are shown in stick. The two deletion variants (pink and gold) could disrupt all of the interactions by shortening or unwinding the helix. **g**, PMG indels in M7 and M10 that are published (black), and similar indels from public databases (orange). **h**, M10 indels keep the sequence in-frame but add or delete one or more amino acids. The published indels are in the five black amino acids. Similar indels found in databases overlap and also extend further in each direction, as shown in gray. The C-terminal extension of M10 makes a sharp turn and docks in the α subunit. This creates a protrusion that also includes the adjacent L8-9 loop. **i**, Although only two residues in M10 interact with β subunit directly, three residues of the docked C-terminus interact with six residues in β. Consequently, changes to the length of M10 should have profound effects.

A completely different pathogenic mechanism is suggested by other ATP1A3 PMG variants. There are five PMG missense variants (yellow) in the L7-8 extracellular loop (**Fig. 3a**)^13-15^. Four are in amino acids at the extracellular interface of α and β subunits, and they contact amino acids in the β subunit headpiece. The fifth does not contact β directly but crucially underpins the structure of the L7-8 loop (purple). This loop is as essential for α-β assembly as β’s transmembrane span^44^. There is also a splice variant in the L9-10 extracellular loop where lysine normally interacts with two β subunit residues (**Fig. 3d)**^14^. Mis-splicing there should also weaken the interaction with β.

Where else in the structure do β and α associate? **Fig. 3e** shows all of the β subunit residues that interact with α in spacefill; the yellow ones are in contact with PMG variants or perturbed by deletions in M7, and the white ones are the rest. M7 has extensive interactions with β’s transmembrane span (**Fig. 3f**). Only one PMG missense variant has been reported there^14^, but more significantly, there are two PMG-associated deletions^13,15^: a single residue (one case) or three adjacent residues (3 cases, different DNA sequences). Because the deletions will shorten, rotate or unwind M7, they could perturb all of M7’s interactions with β’s span. Interestingly, three more M7 indel variants are currently found in ClinVar without phenotype information (**Fig. 3g**).

A cluster of PMG indel variants in M10 draws attention to a structure where a protein could conceivably dock. There are PMG-associated in-frame deletions and insertions (indels) near the cytoplasmic end of M10 that are remarkably prevalent for a short stretch of amino acids (**Fig. 3h**). Four papers reported indels in seven patients with documented PMG^13,15,18,45^. Seven similar indels are found in ClinVar, Decipher, and LOVD (**Fig. 3g**). Although their phenotype information isn’t available, their striking prevalence in databases suggests a clinical bias for detection. The published indels lengthen or shorten M10 by one or two residues in-frame in residues 991 to 995 (**Fig. 3h**). Near the cytoplasmic surface, M10 juts out from the bundle of transmembrane helices, as does the adjacent intracellular loop between M8 and M9, L8-9 (**Fig. 3h**). In contrast, loop L6-7 extends into the cytoplasm interior, behind the bulge, creating space. At the end of the M10 helix, the C-terminus normally docks into the protein interior, but this should be impaired by the M10 length changes. This raises the question as to whether the M10-L8-9 bulge—or the undocked C-terminus—might have a different role. The M10 bulge and adjacent loop L8-9 are at the polar edge of the lipid bilayer, as defined by resolved phospholipids in Na,K-ATPase structures, and they do not change with conformation during ion transport. This leaves three possibilities. One is that loosening of α-β interactions in any one of three locations, extracellular, membrane, or cytoplasmic, is pathogenic at a specific step in neurodevelopment. Miyatake et al. 2021 hypothesized that PMG variants may prevent the association of β with α, and they obtained some promising but inconsistent evidence^13^. It would be a novel form of pathogenicity for Na,K-ATPase, but it seems mechanistically unlikely because association with β is required for α to exit from endoplasmic reticulum^2^.

### Protein-disrupting variants (LoF)

Protein-disrupting variants (termed loss of function in genetics) include large deletions, stop gain or stop loss, frame shifts, and splice site variants that are predicted to induce complete abolition of protein function. ATP1A3 has a loss-of-function (LoF) observed/ expected upper bound fraction (LOEUF) score of 0.2, indicating low tolerance for LoF variation^3^. However, clinical results in ATP1A3 cases show that pathogenicity varies greatly (**Fig. 4**).^14,46-49^ There are also two large deletions. One deletes the 2^nd^-16^th^ exons (out of 23) where the individual was normal at birth and developed seizures that may result from a different variant in GLUT1 (unpublished). The second deletes everything from exon 21 through the 3’UTR, and the individual has severe disease, DEE, per ClinVar.

**Figure 4.**
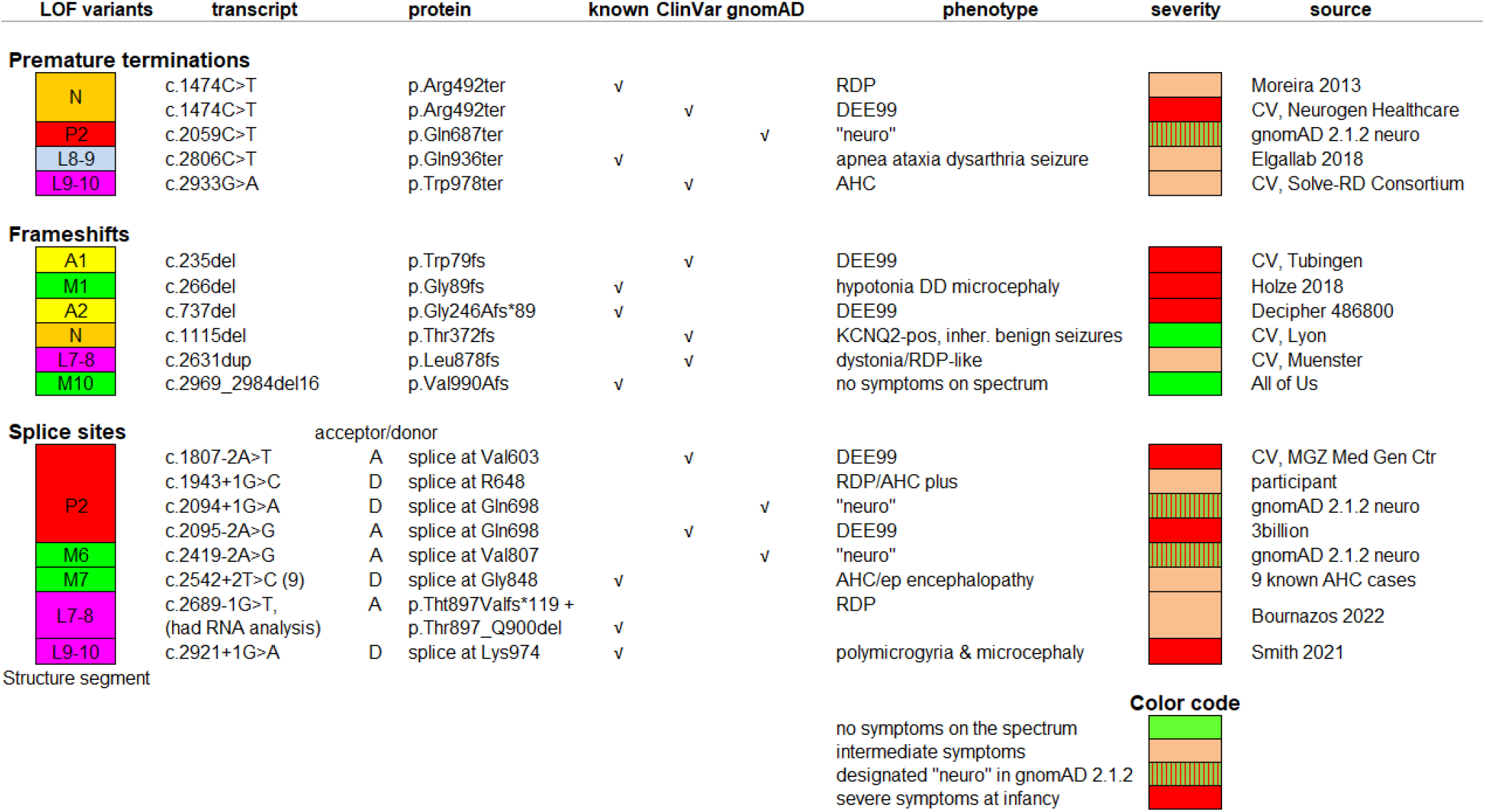
LoF variants are associated with a wide range of clinical phenotypes, even asymptomatic. Premature terminations, frameshifts and splice sites are all considered LoF because they cannot produce the normal protein. Nonetheless they can be pathogenic and dominant. If the aberrant mRNA is not degraded by nonsense-mediated decay, the protein may be degraded by ER-associated degradation by proteasomes. However, the presence of translated nonsense sequence that cannot fold or aggregates may activate the ISR or UPR and even result in apoptosis. The efficacy of these defensive processes may vary considerably. The column of color blocks on the left match the colors in structure images except that here transmembrane spans are colored green. Each line is one case except for multiples of the splice variant at c.2542. Variants are identified as known (published or not), found in ClinVar with phenotype indications (CV), or previously recorded in gnomAD 2.1.2 as “neuro”.

We can postulate that the different clinical results are because aberrant mRNA is effectively eliminated in the first case by nonsense-mediated decay (NMD), but in the second case, which is not subject to NMD because the final exon is deleted, mRNA is translated to protein with a toxic consequence such as misfolding and protein aggregation^50^.

The pathogenicity of a splice site variant additionally depends on whether it is in the donor or acceptor splice sequence because any cryptic splice sites that might rescue it will be different. For the splice site at Lys974, a donor variant produced PMG and microcephaly, while an acceptor variant was apparently benign (**Fig. 4**). Also, there are nine AHC cases for donors at the c.2542 splice site (three different nucleotide changes) but no acceptor cases at c.2543. Experimental investigation of consequences in mRNA, both for sequence abnormality and for abundance relative to WT, is needed in such cases^49^.

A non-binary outcome is found in many other protein-disrupting variants (**Fig. 4**). There are individuals that are symptom-free (2 cases); severely affected (7 cases); designated “neuro” in gnomAD 3.1.2 but where severity is unknown (3 cases); and there are 16 individuals with intermediate symptoms on the ATP1A3 spectrum, including 7 singleton cases and 9 AHC cases at the splice site. This heterogeneity of phenotypes is likely due to haploinsufficiency when mRNA decay is effective, and to variability in the degradation rate or the toxicity of aberrant protein products. There are currently 24 additional protein-disrupting variants in ClinVar and 22 in gnomAD 4.1.

## Discussion

Pathogenicity due to loss of activity, gain of aberrant function, and maladaptive protein misfolding during biosynthesis applies to proteins broadly. The structural correlations we observe entail different mechanisms and several types of protein secondary structure.

The variants with the mildest syndromes were absent from the ion pathway. In the P domain they were in its peripheral helices and not in its central β-sheet. Prior in-depth investigations of RDP variants have uncovered functional alterations in activity or ion affinity^38^; lower affinity results in less activity in physiological conditions^22^. It is apparent that these affinity differences are due to long-range effects of their local perturbations.

Variants causing AHC and cerebral palsy-like spasticity clustered near the ion pathway, likely altering ion binding or movement, a gain of toxic function^12^. Experiments employing electrophysiology have uncovered a variety of ways that mutations of Na,K-ATPase produce aberrant flow of either sodium or protons or alter the 3:2 stoichiometry^51-53^. Variants that affect the ion pathway or perturb hydrogen bond networks in the stalk can cause leaks^12,27,54,55^.

The third category of pathogenicity entails activation of the UPR or ISR to combat misfolded protein^40,56^. For this kind of pathogenicity, abundance matters. Basal quality control mechanisms may dispose of low-density misfolded proteins easily, but Na,K-ATPase is abundant. Neurons may redirect so many resources to the ISR or UPR that growth, differentiation, and maintenance are impaired, or they may die. Here we saw that DEE99 variants are divided into two groups based on location. In the membrane domain they often replaced glycines or altered prolines, the amino acids that make tight turns possible or break helices. In the P domain, DEE99 variants in the core β-sheet should prevent P1 and P2 from interdigitating, preventing domain assembly and causing folding failure^57^. Both routes result in a protein the cell cannot finish, consistent with the misfolding and stress responses reported for variants of this class.

The fourth category of pathogenicity emerged from examining the structural locations of polymicrogyria variants. Three were similar to DEE variants in the transmembrane domain, not surprising considering their symptom overlap. Many PMG variants, however, were in locations unrelated to transport or catalysis, at the interface with β subunit or as indels in the C-terminal bulge.

Aberrant RNA transcripts (splice site variants, premature terminations, frame shifts and large deletions) all have the potential to produce frameshift nonsense or just incomplete sequence that will remain unfolded. When the cell makes a protein that it cannot complete, it has pathways to prevent its translation or degrade the protein product. The efficacy of these defenses will determine the clinical outcome. The LoF variants in this gene illustrate the challenge of determining whether LoF variations are pathogenic or not.

In conclusion, the clinical spectrum of this gene points to vulnerable structural elements that have universality. The way phenotype severity of ATP1A3 variants correlates with problems of protein folding calls for incorporation of vulnerable features into pathogenicity prediction. By extension, the uncertainty about pathogenicity of aberrant LoF transcripts applies to all variants that might impair protein folding and elicit cellular defensive mechanisms.

## Methods

The atomic structures used were of *Sus scrofa* ATP1A1 (α subunit) in complex with ATP1B1 (β subunit) and FXYD2 (γ, a small modulator). The paralogs ATP1A1, ATP1A2 and ATP1A3 are 88-89% identical. Among the paralogs there are many clinical cases in corresponding amino acids: ATP1A3 variants in 47 amino acids have occurred in one or both of ATP1A1 and ATP1A2. Use of high resolution ATP1A1 structures is justified here because differences between the paralogs are in clusters at the protein surface, and none of the ATP1A3 clinical variants were in paralog-specific amino acids. The high resolution structures we examined encompass multiple conformations from different steps of the ion transport cycle: PDB files 3WGU (E1∼P.ADP.3Na^+^)^58^, 8JBK (E1.3Na^+^)^59^, 3KDP (E2.2Rb^+^)^60^, 7Y45 (shark E2.2K)^61^, and 7WYT (shark E2.ouabain)^62^ from X-ray crystallography or cryo-electron microscopy. Recently new cryoEM structures of purified ATP1A3 have been published with some additional intermediate conformations that could provide more insights^63^.

Figures used 3WGU unless otherwise stated because it has the complete structure of the Na,K-ATPase β subunit N-terminus. Structures are shown in ribbon view, with spacefill of the backbone atoms to show the positions of amino acids that have variants, and sometimes with individual amino acids in stick view to visualize their sidechains or spacefill to illustrate shape. Swiss PDB Viewer V4.2 was used for visualization, measurements of distances between atoms, rotamer manipulation, detection of hydrogen bonds, detection of clashes, and superposition and alignments of structures being compared. Van der Waal contacts are in the range of 3.3-4.5Å, and we arbitrarily defined contact between amino acids as being 4Å or less. In the literature, residue numbering varies both because of species and paralog differences and because of a precedent that amino acid numbering in ATP1A1 should begin with the first residue after the known cleavage of five amino acids at the N-terminus. Residue numbering here is according to the 1013 amino acid human ATP1A3 mRNA transcript, NM_152296.5, in line with clinical usage.

## Acknowledgements

The authors acknowledge the foundational work of the many investigators who have determined Na,K-ATPase structure and mechanism, and those who have carried out exacting analysis of Na,K-ATPase mutations by biochemistry and electrophysiology. We thank the ATP1A3 patient organizations for fostering scientific advances by sponsoring annual international meetings. This work was supported by the National Institutes of Health – National Institute of Neurological Disorders and Stroke (R01NS058949), and by the Cure AHC Foundation, the Alternating Hemiplegia of Childhood Foundation, and RARE Hope.

